# Constructing the hierarchy of predictive auditory sequences in the marmoset brain

**DOI:** 10.1101/2021.10.25.465732

**Authors:** Yuwei Jiang, Misako Komatsu, Yuyan Chen, Ruoying Xie, Kaiwei Zhang, Ying Xia, Peng Gui, Zhifeng Liang, Liping Wang

## Abstract

Our brains constantly generate predictions of sensory input that are compared with actual inputs, propagate the prediction-errors through a hierarchy of brain regions, and subsequently update the internal predictions of the world. However, the essential feature of predictive coding, the notion of hierarchical depth and its neural mechanisms, remains largely unexplored. Here, we investigated the hierarchical depth of predictive auditory processing by combining functional magnetic resonance imaging (fMRI) and high-density whole-brain electrocorticography (ECoG) in marmoset monkeys during an auditory local-global paradigm in which the temporal regularities of the stimuli were designed at two hierarchical levels. The prediction-errors and prediction updates were examined as neural responses to auditory mismatches and omissions. Using fMRI, we identified a hierarchical gradient along the auditory pathway: midbrain and sensory regions represented local, short-time-scale predictive processing followed by associative auditory regions, whereas anterior temporal and prefrontal areas represented global, long-time-scale sequence processing. The complementary ECoG recordings confirmed the activations at cortical surface areas and further differentiated the signals of prediction-error and update, which were transmitted via putatively bottom-up γ and top-down β oscillations, respectively. Furthermore, omission responses caused by absence of input, reflecting solely the two levels of prediction signals that are unique to the hierarchical predictive coding framework, demonstrated the hierarchical predictions in the auditory, temporal, and prefrontal areas. Thus, our findings support the hierarchical predictive coding framework, and outline how neural circuits and spatiotemporal dynamics are used to represent and arrange a hierarchical structure of auditory sequences in the marmoset brain.

## Introduction

The mammalian cerebral cortex is organized as a functional hierarchy, through which information propagates from the lowest levels of primary sensory or motor cortices to the higher levels (Fuster, 1997; Huntenburg et al., 2018; Mesulam, 1998). At each processing level, neurons integrate information from multiple neurons at the level below, thus encoding increasingly abstract information over ever-larger temporal and spatial scales. Meanwhile, the reciprocal connections between cortical areas provide neurons with feedback from the level above (Felleman & Van Essen, 1991; Marques et al., 2018). In the time domain, the processing hierarchy is particularly important for cognition requiring a temporal integration of sensory stimuli and on-line action planning, such as spatial navigation and language (Dehaene et al., 2015; Giraud & Arnal, 2018).

The hierarchical predictive coding theory offers a unified framework for this functional hierarchy. It states that the brain develops a generative model of the world that constantly predicts sensory input (Bizley & Cohen, 2013; Rao & Ballard, 1999; Spratling, 2010). The comparison of predicted and actual sensory input then updates an internal representation of the world (Keller & Mrsic-Flogel, 2018). This process occurs throughout the cortical hierarchy. Specifically, the higher levels of the hierarchy send a top-down signal to lower levels in the form of a prediction of the bottom-up input to that area. The difference computed between the prediction and the bottom-up input is the prediction-error. The key assumption of the theory is that the prediction-error propagates across different depths of hierarchy, and in turn, the prediction propagates backward, providing signals to update the internal model at each level. Much of the research focuses on the signals of prediction and prediction-error at one unique level, such as the level of spatial detection in visual cortex (Attinger et al., 2017; Fiser et al., 2016) or one scale of temporal expectations in auditory cortex (Gagnepain et al., 2012; Rubin et al., 2016). However, the essential feature of predictive coding, i.e., the notion of hierarchical depth (across multiple levels), is less well investigated (Chao et al., 2018; Parras et al., 2017).

Here, we combined whole-brain 9.4 T functional magnetic resonance imaging (fMRI) and large-scale electrocorticography (ECoG) recordings to gain both spatial and temporal neural information during a hierarchical local-global auditory sequence task in the brains of marmosets (Fig. 1A). These nonhuman primates are an increasingly important animal model for auditory processing because their social behavior and cognition are similar to those of humans (Eliades & Miller, 2017; Miller et al., 2016). It is difficult to record neural activities (both low-frequency event-related potentials and high-frequency potentials, e.g., high gamma band) in the primary auditory cortex in humans and macaque monkeys because of the cortical sulci. As a result, studies on predictive processing in audition are restricted to electroencephalography (EEG) or ECoG recordings in associative cortical areas. By contrast, the smooth surface of the lissencephalic marmoset brain provides a unique opportunity to compare whole-brain fMRI and large-scale surface ECoG signals. In the present study, we used these combined techniques to study the distribution of activities over cortical areas along the auditory pathway. We employed a local-global paradigm enabling us to measure predictive processing at one level along with the conditional propagation of prediction-errors to the next level, depending on the predictive context (Fig. 1B). Both the blood-oxygen-level-dependent (BOLD) signals and the electrophysiological power changes were investigated across a broad frequency spectrum at the whole-brain level. The aims of the present study were to assess the hierarchical depth of predictive auditory sequences and to identify the neural circuitries and temporal dynamics for processing prediction-errors and prediction updates at two distinct levels: the lower local level with a short timescale (i.e., 150 ms) and the higher global level with a longer timescale (i.e., seconds).

**Figure 1:**
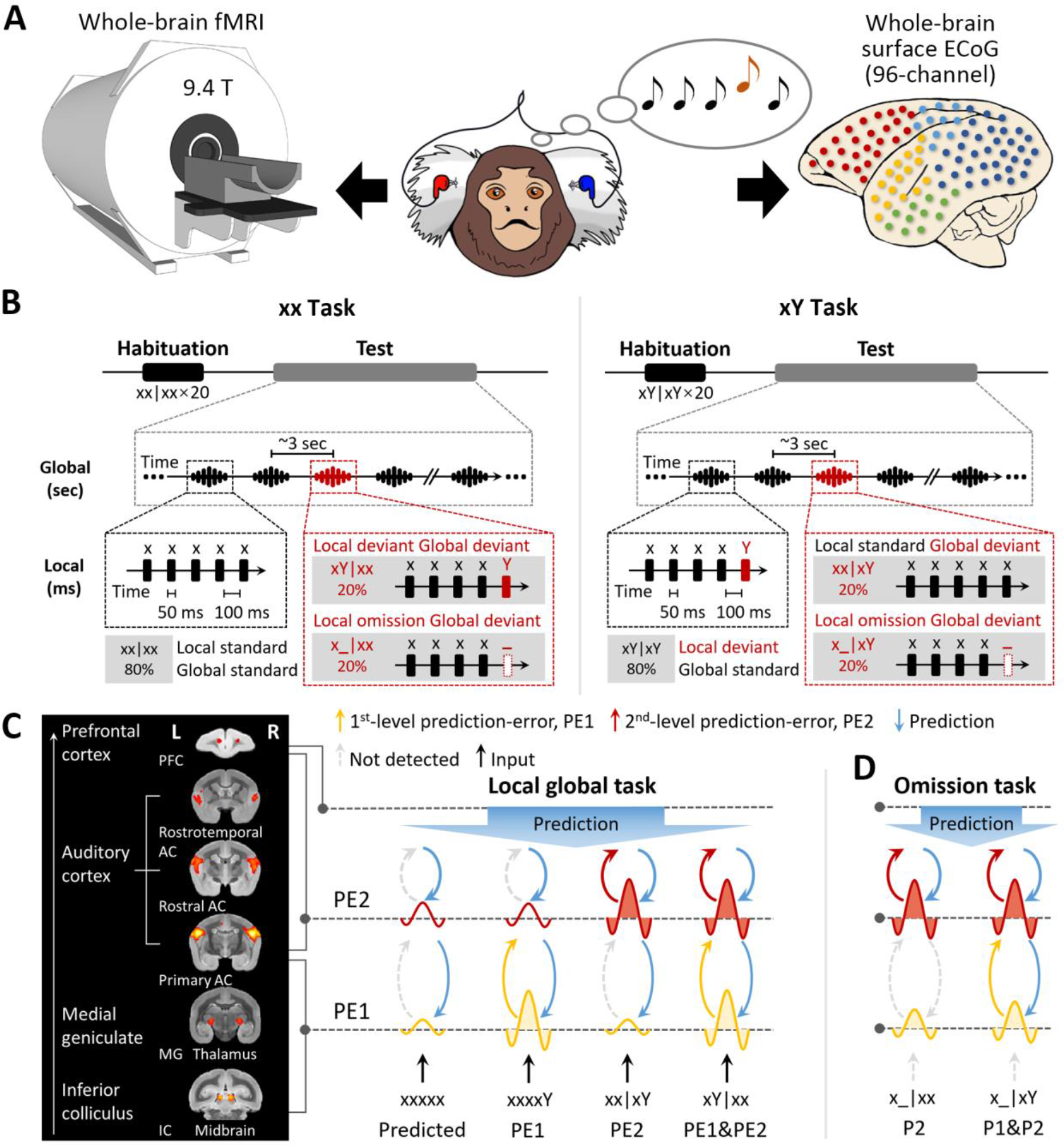
Experimental setup and local-global paradigm. **(A)** Experimental setup of whole-brain fMRI (9.4T) and ECoG (96-channel); awake marmosets passively listen to auditory stimuli. **(B)** Schematic of local-global paradigm (for details, see Materials and Methods). **(C)** *Left*, fMRI activations for auditory sounds relative to baseline, along the auditory pathway, in marmosets. *Right*, hypothesized predictive processing model of the local-global paradigm, corresponding to the auditory pathway. **(D)** Hypothesized predictive processing model of the local-global omission paradigm. P1, 1^st^-level prediction; P2, 2^nd^-level prediction.

## Results

### Local-global paradigm and a two-level hierarchical predictive processing model

We adopted the local-global oddball paradigm (Fig. 1B) (Bekinschtein et al., 2009), in which the auditory sequence comprised five identical tones (“xxxxx”) or four identical tones followed by a distinct tone (“xxxxY” where x and Y can be tones of 800 or 6,000 Hz, alternatively), which are referred to here as “xx” and “xY” sequences, respectively. During an initial habituation phase, five marmosets (three for the fMRI and two for the ECoG experiment) passively heard one sequence type (e.g., xY) in a given block. Then, during the following test phase, we probed for brain responses to novel sequences that either respected the habituated sequence (80% of trials) or violated it (20% of trials; alternative/deviant sequence [xx in this example]). In experiments assessing the effect of an omission, the 5^th^ tone was simply omitted in the violation sequence (comprising 20% of trials) (Fig. 1B; for details, see Materials and Methods).

We used this paradigm to test the hypothesis of hierarchical neural dynamics during auditory sequence processing (Fig. 1C), which predicts how the brain informs estimated statistics on the basis of the history of auditory sequences at two levels.

1. At the 1^st^ (“local”) level, expectation violations result from tone-to-tone transition probability, which uses the most recent (150 ms) observations only, e.g., the perception of an oddball (Y) tone deviating from four repetitions of standard (x) tones. The pattern of brain activation of predicted violation at this level is shown in Fig. 1C (in yellow), where the Y tone induces significantly greater activation than the x tone in both xx and xY tasks, referred to as the 1^st^-level prediction-error (PE1).
2. These local sequences, whether normal (xx) or oddball (xY), are repetitively presented to form 2^nd^-order sequences (sequence to sequence), thereby generating expectations at a 2^nd^ (“global”) level, which takes into account all five tones (∼3 s). The 2 × 2 experimental design enables us to probe the neural responses to two types of global-level violations.

a. Violation only at the 2^nd^ level: an expectation violation results from the perception of the xx sequence deviating from the xY standard sequence (xx sequences in xY task, referred to as xx|xY). As shown in Fig. 1C (in red), the xx violation sequences should induce significantly higher activation than the xY standard sequences, but they do not produce higher activation with the xx sequence at the local level. Here, the violation xx at the 2^nd^ level is referred to as PE2.
b. Violation at both 1^st^ and 2^nd^ levels: expectation violations resulting from the perception of the xY sequence deviating from the xx standard sequence (xY sequences in xx task, referred to as xY|xx) should generate two successive violations. Because the two levels of violations are activated sequentially, the 1^st^-level novelty (i.e., the 5^th^ tone is Y) determines and triggers the 2^nd^-level novelty (the xY sequence), referred to as a PE1&2 violation (Fig. 1C).
3. The brain produces predictions at both local tone (Y) and global sequence (xx or xY) levels. When the incoming signal is omitted, brain responses should reflect solely the prediction signals and how they varied depending on the current context (xx or xY task) (Fig. 1D). Specifically, in the xY task, where two successive oddball predictions are generated, we should observe a neural response to omission at both the 1^st^ and 2^nd^ levels, corresponding to the predictions of the Y tone and of the xY sequence. In the xx task, however, without the prediction of oddball tone in the xx sequence, the response to omission of the x tone may not be as strong as that of the Y tone at the 1^st^ level, where only the prediction of the xx sequence at the 2^nd^ level should be as robust as that in the xY task.

To test these hypotheses, we combined 9.4 T fMRI and 96-channel whole-brain ECoG of awake marmosets to assess the whole-brain neural circuits and temporal dynamics during hierarchical auditory sequence processing, and searched for the neural representations of prediction-errors and predictions at two different levels.

### 1st-level (local) novelty (xY sequences)

We first examined our data for the presence of a local mismatch response evoked by the deviance of the 5^th^ tone. fMRI revealed the cortical regions activated by the novel tone, defined by the conjunction analysis using the contrasts between xY- and xx-evoked responses in both xx and xY tasks: (xY|xx – xx|xx) ∩ (xY|xY – xx|xY) (see Materials and Methods). In the three monkeys, significant activation was found bilaterally along the auditory pathway, from the inferior colliculus (IC) in the midbrain and medial geniculate nucleus (MG) in the thalamus to auditory cortex (including auditory core and belt areas), the temporoparietal transitional area, dorsal prefrontal cortex, and anterior/posterior cingulate cortex (Fig. 2A, red-yellow; *p* < 0.05, false-discovery rate [FDR] corrected; Table S1; individual monkeys: Fig. S1A). These areas showed positive activation (relative to the mean of the baseline in this run) for each auditory sequence and significantly higher activation for the Y tone in any xY sequence, regardless of the standard sequence pattern (xx or xY).

**Figure 2:**
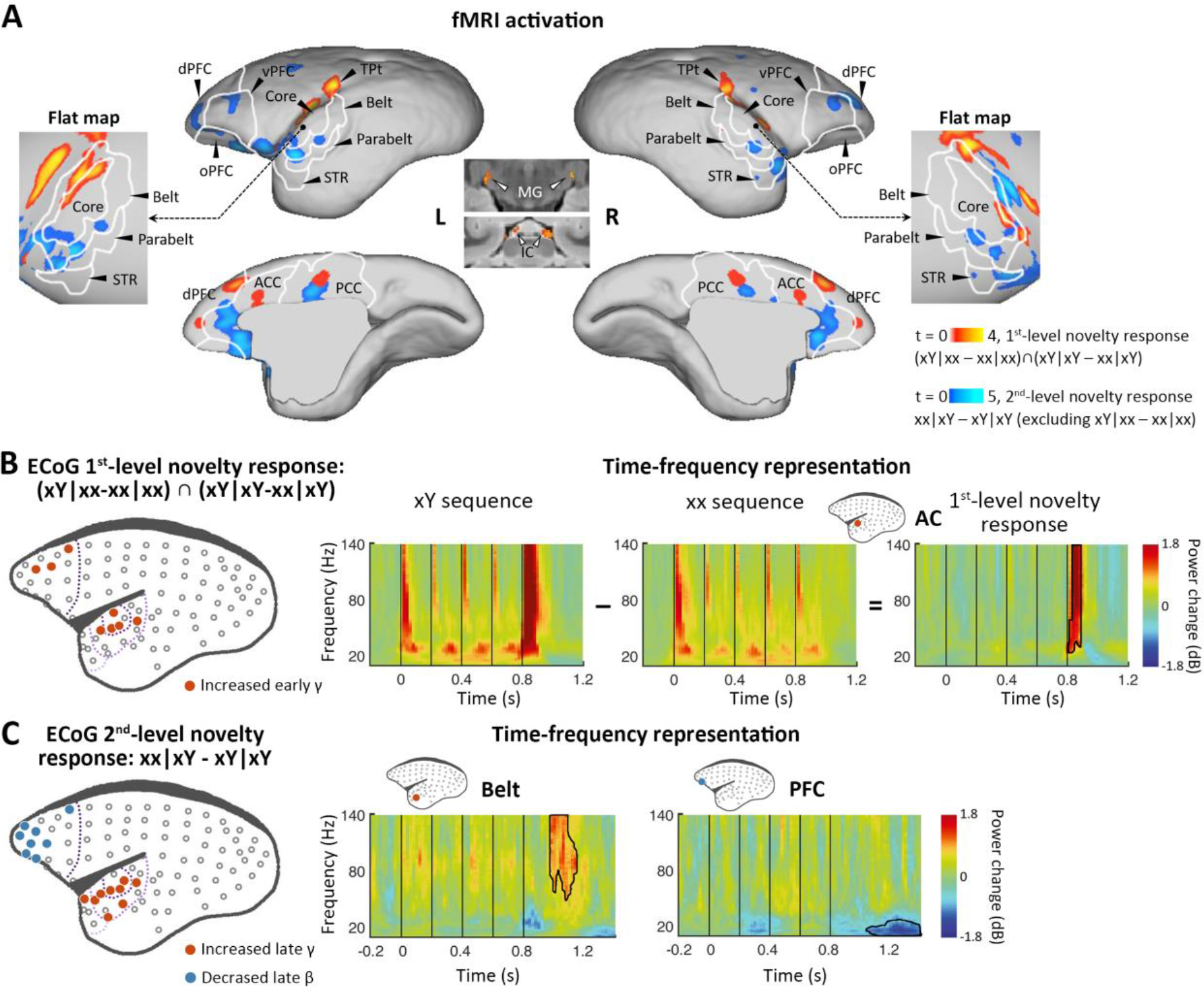
Neural activations to the 1^st^-level local and 2^nd^-level global novelties, respectively. **(A)** *Red-yellow*, fMRI activations to 1^st^-level novelty identified by subtraction in both xx and xY tasks; *blue-light blue*, fMRI activations to the 2^nd^-level novelty identified by subtraction only in the xY task (*p* < 0.05, FDR-corrected). L, left; R, right; IC, inferior colliculus; MG, medial geniculate nucleus; STR, superior temporal rostral area; dPFC, dorsal prefrontal cortex; vPFC, ventral PFC; oPFC, orbital PFC; ACC, anterior cingulate cortex; PCC, posterior cingulate cortex; TPt, temporoparietal transitional area. **(B)** *Left*, Localization of ECoG electrodes for local novelties (monkey J, *p* < 0.05, cluster-corrected). The gray dots represent all 96 electrodes. The dotted lines from dark to light violet indicate frontal, auditory core, auditory belt/parabelt, and STR. *Right*, Time-frequency representation (TFR) of representative AC electrodes with 1^st^-level novelty response, generated by comparing local deviants (xY sequence) with local standards (xx sequence). The vertical lines indicate the onset times of five tones in each sequence. **(C)** *Left*, Localization of ECoG electrodes for global novelties (monkey J, *p* < 0.05, cluster-corrected). *Right*, TFR of representative electrodes in the auditory belt cortex and PFC.

Because of the relatively poor temporal resolution of BOLD signals and the lack of time- frequency information with fMRI, as a complement, we performed ECoG (96-channel) recordings with the same auditory paradigm in two additional marmosets. The ECoG system acquires high-fidelity broadband neural signals from an entire cortical hemisphere, with balanced spatial, spectral, and temporal resolutions. The spatial, spectral, and temporal dynamics of ECoG signals were quantified by the time-frequency representation (TFR) (see Materials and Methods). The 1^st^-level novelty response was defined as a significant difference in TFRs for the 5^th^ tone (between Y and x) in both xx and xY tasks. Consistent with the regional activation observed in the fMRI experiment, the electrodes (Fig. 2B and Fig. S2A, electrodes in red; *p* < 0.05, cluster-corrected) exhibited an increase in γ-band power (>40 Hz) right after the 5^th^ tone (peak at 74 ± 34 ms after the 5^th^-tone onset) (Fig. 2B and Fig. S2A, right). In both marmosets, these electrodes responding to 1^st^-level novelty also displayed the highest responses to auditory sounds relative to the baseline (Fig. S3; *p* < 0.05, cluster-corrected), suggesting that the early γ-band power increase was most likely from the primary auditory cortex.

### 2nd-level (global) novelty (xx sequences in the xY task [xx|xY])

We next looked for a 2^nd^-level novelty response dependent on the overall sequence rather than individual tones. At this level, two internal sequence representations (xx and xY) were established during the habituation period, which enabled us to test the two different types of global novelties. We first tested the novelty responses in the xY task, where the deviant sequence (xx) only generated a violation at the 2^nd^ level because it did not include local novelty. We then used fMRI to identify the brain regions showing significantly higher activation for xx sequences than for xY sequences in the xY task, and no difference between the xx and xY sequences in the xx task (i.e., when the sequence suddenly changed from xY to xx, but not from xx to xY). Unlike what was observed with the 1^st^-level novelty, we identified higher-order regions along the auditory pathway, especially in the temporal-prefrontal pathway and anterior-posterior cingulate areas, including bilateral anterolateral auditory area, superior temporal rostral area (STR), and prefrontal cortex (Fig. 2A, blue-light blue; *p* < 0.05, FDR-corrected; Table S2; individual marmosets: Fig. S1B).

ECoG showed that the 2^nd^-level novelty generated a later γ-band power (peak at 242 ± 17 ms) after the 5^th^-tone onset than observed for the 1^st^-level novelty (Fig. 2C and Fig. S2B, electrodes in red; *p* < 0.05, cluster-corrected). The extended γ-band power was from the electrodes located in the anterior auditory and superior temporal cortices. Furthermore, the frontal electrodes, in regions similar to those identified in the fMRI experiment, showed a late β-band power decrease (12–30 Hz) starting at 206 ± 170 ms after the 5^th^-tone onset, with a longer latency and lasting for more than 300 ms (Fig. 2C and Fig. S2B, electrodes in blue; *p* < 0.05, cluster-corrected).

Since the γ and β bands are thought to subserve bottom-up and top-down communications, respectively, in both humans (Arnal & Giraud, 2012; Michalareas et al., 2016) and monkeys (Bastos et al., 2015; van Kerkoerle et al., 2014), the ECoG results confirmed the activated regions shown by fMRI and, more importantly, functionally dissociated the fMRI activation patterns between frontal and temporal regions (Fig. 2A). Combined with the finding of increased early γ-band activity shown in the 1^st^-level novelty response, we thus propose that the early and late increases in γ-band activity in the auditory and temporal cortices are associated with the bottom-up prediction-errors PE1 and PE2, respectively, and that the long-lasting decrease in β-band activity in frontal areas likely represents the subsequent top-down prediction or updating process.

### Violation at both 1^st^ and 2^nd^ levels (xY sequences in the xx task [xY|xx])

We next examined the violation sequence xY in the xx task, which generates novelty responses at two successive levels. In this analysis, the internal representation is for the xx sequence; thus, the conjunction analysis [(xY|xx > xx|xx) ∩ (xY|xY = xx|xY)] was used.

That is, we searched for brain regions showing significantly higher activation for xY sequences than xx sequences in the xx task, and no difference between xx and xY sequences in the xY task. As predicted, the xY sequence produced a violation at both the 1^st^ and 2^nd^ levels, as the strongest activation during fMRI encompassed the low-level auditory pathway, including the MG, auditory core and belt regions, as well as the parabelt region of the higher-order auditory cortex and progressing to the temporal-frontal network, e.g., STR, A8, and A10 (Fig. 3A; *p* < 0.01, FDR-corrected; Table S3; individual marmosets: Fig. S1C).

**Figure 3:**
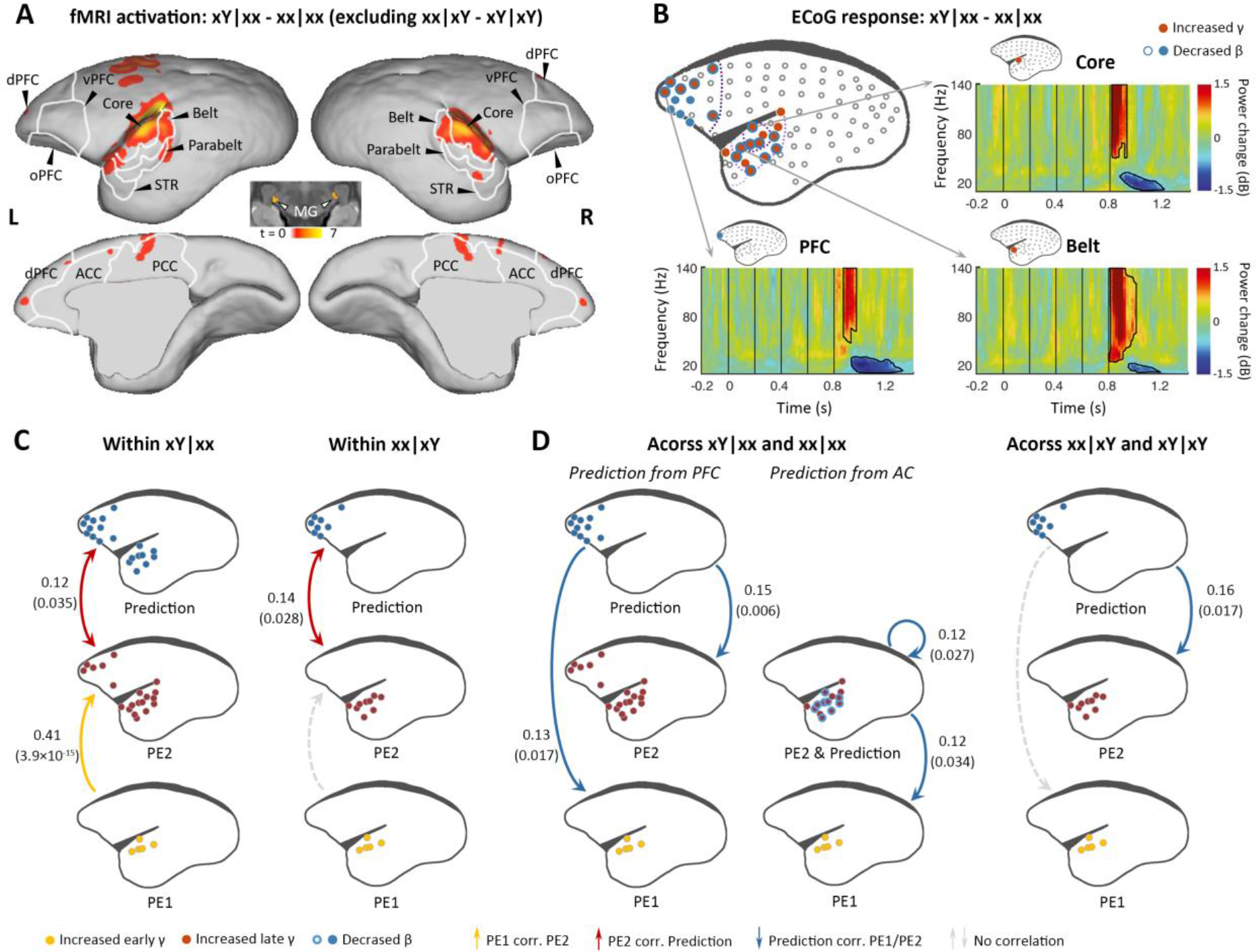
Brain activations to the two successive (local and global) novelties. **(A)** fMRI activations to both local and global novelties (*p* < 0.01, FDR-corrected). **(B)** Significant ECoG responses detected by xY minus xx sequences in the xx task (monkey J, *p* < 0.05, cluster-corrected). **(C)** Diagram of the functional correlation among 1^st^- and 2^nd^-level prediction-errors (PE1 and PE2, respectively) and prediction signals within deviants in the xx task (left, xY|xx) and xY task (right, xx|xY) (monkey J). **(D)** *Left*, diagram of the functional correlation across trials, showing the correlation between prediction signals on xY|xx and PE1/PE2 signals on subsequent standards (xx|xx directly appeared after xY|xx). *Right*, the functional correlation across xx|xY and the subsequent xY|xY trials (monkey J). The dots in the diagrams indicate the electrodes with significant responses that used in the correlation test. Lines represent significant correlations, while arrows indicate temporal orders of the signals. Labeled values are in forms of Pearson correlation coefficient (*p*- value).

ECoG confirmed the activations along the auditory and temporal-frontal pathways, and most of the electrodes located at the surface of the superior temporal gyrus showed the early γ-band power increase, possibly denoting the representation of 1^st^-level prediction-error (Fig. 3B and Fig. S2C; *p* < 0.05, cluster-corrected). The detection of the change from tone x to Y signals a violation of sequence, and accordingly, the extended γ-band power increases (early and late γ-band power increases jointly lasting >200 ms) occurred in the anterior auditory and superior temporal regions (Fig. 3B and Fig. S2C, electrodes in red; *p* < 0.05, cluster- corrected). In the marmoset (J), the increased γ-band power also occurred for the prefrontal electrodes, suggesting a propagation of error signals from the 1^st^ to the hierarchically higher 2^nd^ level. Furthermore, both the frontal and auditory electrodes revealed decreases of the long-lasting β-band power (starting at 148 ± 37 ms after the 5^th^-tone onset and lasting >300 ms) (Fig. 3B and Fig. S2C, electrodes in blue; *p* < 0.05 cluster-corrected). Thus, these data suggest a potential two-level top-down modulation process of tones and sequences.

### Functional correlation between γ- and β-band power within and across trials

The proposed PE1/PE2 and prediction update represented by the early/late γ-band and long-lasting β-band power, respectively, were based on time-frequency characteristics. To verify a functional relationship between the frequency band and predictive function, we examined the directional correlation of corticocortical interaction in the novelty responses using the observed signals of PEs (γ-band activity) and prediction/updating (β-band activity) within and across trials (see Materials and Methods). As shown in Fig. 1C, the current framework predicts that the activation of the 1^st^-level violation should determine the activation at the 2^nd^ level, especially on xY trials in the xx task. Thus, within a deviant trial, the hypothesis is that the early γ-band power increase (1^st^-level error) will trigger the late γ-band power increase (2^nd^-level error) and then interact with the late long-lasting β-band power decrease (prediction updating). Across trials, the β-band activity should update the internal representations of tones and sequences, which will affect the processing of subsequent trials. In this way, trial-by-trial fluctuations in the β-band power will affect the top-down predictions, and the amount of change will affect prediction-errors on subsequent trials.

We first examined the correlations between the three signals (early γ-band, late γ-band, and late β-band power) within the deviant trial xY|xx. Early γ-band (from primary auditory cortex, starting at 40 ms [J] and 60 ms [N] after the 5^th^-tone onset) and late γ-band (from auditory and superior temporal cortices, starting at 120 ms [J] and 160 ms [N] after the 5th-tone onset) prediction-error signals were positively correlated (Fig. 3C and Fig. S4A, left yellow), and late γ-band prediction-error and β-band (from auditory and prefrontal regions, starting at 140 ms [J] and 220 ms [N] after the 5th-tone onset) prediction-updating signals were positively correlated (Fig. 3C and Fig. S4A, left red). However, within the deviant trial xx|xY, we only observed a significant correlation between late γ-band and late β-band power (Fig. 3C and Fig. S4A, right) as hypothesized, as the 2^nd^-level error was not induced by the 1^st^-level error in the xY task.

Across trials, we investigated whether the long-lasting β-band signal in a deviant trial would affect the early/late γ-band signals in the subsequent trial, a standard (xx|xx) trial according to the experimental design. The prediction signal (β-band) from frontal cortex correlated significantly with early (PE1) and late (PE2) γ-band signals on the following xx|xx trial (Fig. 3D and Fig. S4B, left). Indeed, we also observed significant correlations between prediction signal from auditory cortex and PE1/PE2 signals from auditory cortex (Fig. 3D and Fig. S4B, left). By contrast, prediction signal from prefrontal cortex only significantly correlated with the late γ-band signal (PE2) from auditory cortex on the following xY|xY trial (Fig. 3D and Fig. S4B, right), suggesting a selective update of PE1 and PE2 based on different task contexts. These results demonstrate the neural sequential processing of the prediction-error signals from PE1 to PE2 (i.e., from primary auditory cortex to high-order auditory and superior temporal regions), which were propagated to frontal cortex where they interacted with prediction signals; in turn, the prediction signals subsequently affected the PE1 and PE2 signals in response to new sensory inputs in the following trial.

### Omission novelties (x_|xx and x_|xY) induce prediction signals dependent on task context

Lastly, we examined omission responses. As proposed, omission of the 5^th^ tone should reveal the brain’s hierarchical predictions, and the observed omission response should vary according to the expectation induced by the overall sequence context (xx or xY). fMRI was performed during sequences with omissions and during standard sequences in the xx or xY task (see Materials and Methods). In the xx task, the omission induced the strongest activation in the posterior cingulate cortex, prefrontal cortex, and inferior temporal (TE3) regions (Fig. 4A, red-yellow; *p* < 0.001, uncorrected; Table S4; individual marmosets: Fig. S5A). However, responses to the omission in the xY task were found in the auditory belt cortex as well as the prefrontal and anterior cingulate cortices (Fig. 4A, blue-light blue; *p* < 0.05, FDR-corrected; Table S5; individual marmosets: Fig. S5B).

**Figure 4:**
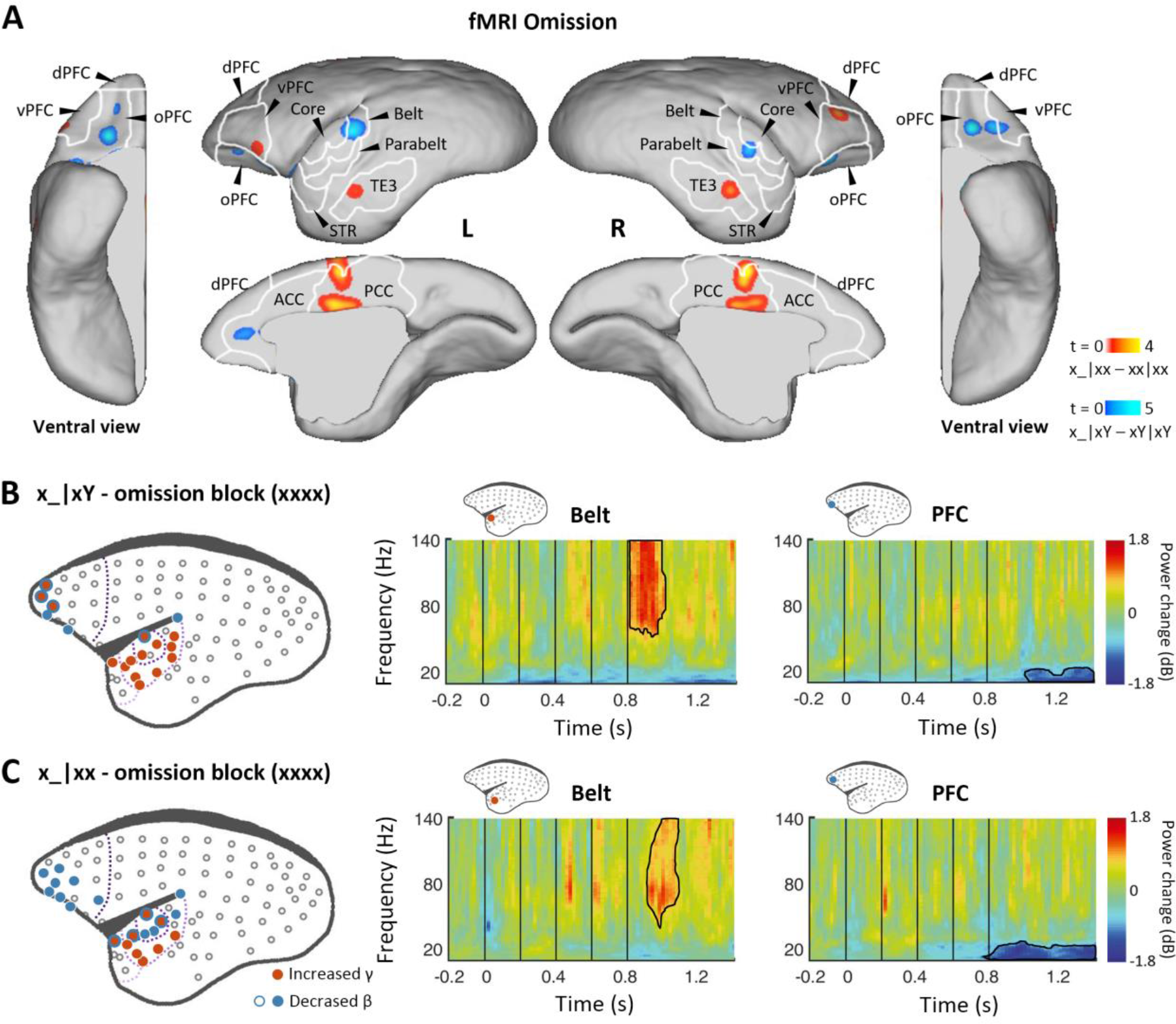
Prediction-error and prediction responses to omission of the 5^th^ tone. **(A)** fMRI revealed areas responsive to an omission of the 5^th^ tone in the xx (red-yellow, *p* < 0.001, uncorrected) and xY tasks (blue-light blue, *p* < 0.05, FDR-corrected). TE3, inferior temporal cortex. **(B)** *Left*, localization of ECoG electrodes with significant responses when comparing rare omissions in the xY task with frequent omissions in the omission block (consisting of xxxx sequences, where an omission is expected, monkey J, *p* < 0.05, cluster-corrected). *Right*, examples of respective electrode response in the auditory belt area and PFC. **(C)** Significant ECoG responses to the omission of the 5^th^ tone when comparing rare omissions in the xx task with frequent omissions in the omission block (monkey J, *p* < 0.05, cluster-corrected), with the same format as for panel B.

In the ECoG experiment, the omission effect was examined by comparing the omission sequence in the xx or xY task with the expected omission stimuli in the xxxx (four identical tones) block (see Materials and Methods). Sequence x_|xY induced an early effect on γ-band power responses from 48 ± 34 ms after the onset of the omitted tone for electrodes in the regions of primary auditory cortex that were similarly activated by the sequence xY|xx (Fig. 4B and Fig. S2D; *p* < 0.05, cluster-corrected). The early latency of this peak response to the omission is consistent with the hypothesis that the response corresponds to an unfulfilled 1^st^- level prediction (Fig. 1D).

In both xx and xY tasks, the omission also produced a significant effect on the γ-band power at a later time, lasting to 282 ± 19 ms (x_|xx, starting at 151 ± 38 ms) and 224 ± 56 ms (x_|xY) after the onset of the omitted tone (Fig. 4C and Fig. S2E; *p* < 0.05, cluster-corrected). The activated electrodes were located at associated auditory and superior temporal areas, which, as predicted, are similar to the activation areas where the 2^nd^-level (global) effect was observed in both xx and xY tasks (Fig. 2C and 3B). Furthermore, in both tasks, the frontal and auditory electrodes revealed consistent long-lasting (>300 ms) decreases in β-band power, suggesting a feedback top-down updating of prediction in both contexts.

Finally, we compared the effect of omission between the xx and xY tasks, testing the unique role of prediction in the hierarchical predictive coding model. An omission was predicted to have a more profound effect on the xY task when a deviant stimulus was expected. In a region where two successive predictions are generated, we should observe a large response to omission, composed of superimposed activations corresponding to the predictions of the “x” tone and the “xY” sequence. By contrast, only one level of predictions should exist in the xx task; thus, the response to omission should be significantly smaller. The difference between omissions is shown in Fig.5. fMRI revealed greater activation in primary and associated auditory and prefrontal cortices during the xY task (Fig. 5A; *p* < 0.05, FDR-corrected; Table S6; individual marmosets: Fig. S5C). ECoG data similarly showed greater activation in the primary auditory and superior temporal cortices in response to the omission during the xY task than during the xx task (Fig. 5B and Fig. S2F; *p* < 0.05, cluster-corrected, peaking at 85 ± 19 ms after the 5^th^-tone onset). The early omission responses in the xY task reveal the PE1 signal generated by top-down prediction to actual auditory input. Indeed, within deviant trials, we observed that the pattern of functional correlations for x_|xx and x_|xY were similar to xx|xY and xY|xx, respectively (Fig. 5C and Fig. S4C). Furthermore, functional correlation analyses across trials showed that in the xx task, the prediction in the x_|xx trial from frontal cortex correlated significantly with the PE2 in the subsequent trial (Fig. 5D and Fig. S4D, left), suggesting that the prediction signal is updated only at the 2^nd^- level sequence process. In the xY task, the prediction in the x_|xY trial from prefrontal cortex correlated significantly with both PE1 and PE2 in the subsequent trial (Fig. 5D and Fig. S4D, right), indicating that both 1^st^- and 2^nd^-level representations were updated by prediction signals from prefrontal cortex, a backward hierarchical top-down process. Moreover, the prediction in the x_|xY trial from auditory cortex only correlated with the PE1 in the subsequent trial (Fig. 5D and Fig. S4D, right), suggesting the 1^st^-level prediction signal from the auditory cortex.

**Figure 5:**
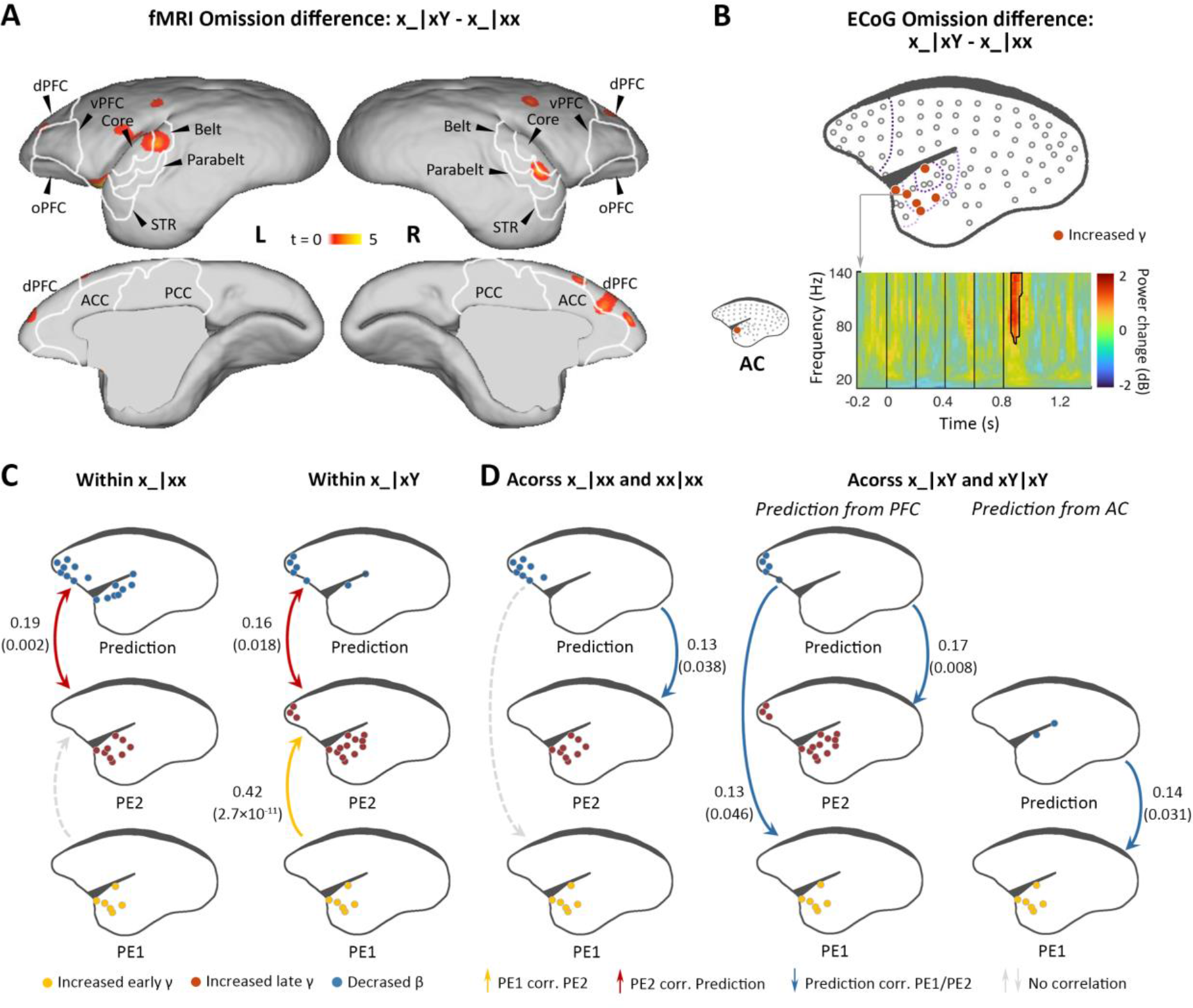
Difference in responses between local deviant omission and local standard omission. **(A)** fMRI revealed areas activated differently between local deviant omission (x_|xY) and local standard omission (x_|xx) (*p* < 0.05, FDR-corrected). **(B)** *Top*, localization of ECoG electrodes with significant differences between x_|xY and x_|xx omissions. *Bottom*, an example of electrode showing significant early γ-band power increases in x_|xY omissions compared with that for x_|xx. **(C, D)** Functional correlation results of x_|xx and x_|xY trials, with the same format as Fig. 3C and D.

## Discussion

We combined 9.4 T fMRI and high-density ECoG with a local-global auditory sequence paradigm to assess the hierarchical depth of auditory sequence processing in both spatial and temporal domains in the marmoset brain. At the whole-brain level, we found progressive encoding of information along the auditory pathway: local auditory information on a short temporal scale is encoded and propagated from the IC and MG, and primary auditory cortex to the posterior and anterior cingulate cortices, where the local 1^st^-level prediction-error (PE1) signal is characterized by the early γ-band. Global auditory information on a long temporal scale propagates from primary auditory cortex to high-order auditory, superior temporal, and prefrontal cortices, where the 2^nd^-level prediction-error (PE2) signals are transmitted via the late γ-band. Furthermore, at both local and global levels, temporal and prefrontal cortices produce prediction feedback signals, which are transmitted in a long- lasting β-band. The prediction signals interact with the PE1 and PE2 signals within a deviant trial as well as with the updated error signals in the subsequent trial. Furthermore, the neural responses to omissions in an auditory sequence at both local and global levels dissected the multiple top-down prediction systems, supporting the hierarchical predictive coding framework.

### Predictive signals in the subcortical areas along auditory pathway

Previous studies in humans and macaque monkeys have suggested that the local novelty could induced neural activations in the primary auditory cortex (Chao et al., 2018; El Karoui et al., 2015; Uhrig et al., 2014; Wacongne et al., 2011). However, due to poor spatial resolution of EEG and compromised signal-to-noise of fMRI (3T), the response to local novelties were barely described in the subcortical regions. Using 9.4 T fMRI in marmosets, we found the neural responses to the local novelty in the IC, MG and primary auditory cortex. Our finding is consistent with a recent mice study with electrophysiological recordings in these regions, where the authors examined local intervals (0.001 – 0.1 s) separating consecutive noise bursts and global rhythmic patterns (∼ 1 s) of inter-burst interval sequences (Asokan et al., 2021). They showed that neural response to local intervals was strong in the IC, but declined across MG and primary auditory cortex; the response to global rhythmic patterns was robust in primary auditory, but weak in MG and IC. Combined with the early peak latency of γ-band power for PE1, the local novelty responses in IC, MG and primary auditory cortex are in line with the thalamocortical correlates of mismatch negativity (Lakatos et al., 2020), suggesting that classical oddball paradigms are involved in local prediction-error. Furthermore, our paradigm with the 2×2 design (e.g., xY|xx novelty) allows to explore the interaction of local and global effect. Interestingly, in the xY|xx condition, the global effect was also observed in MG, suggesting that the auditory regions in the subcortical regions may also process the large timescale auditory information. Future studies using single or multiple units recording would help to clarify the function of sequence processing in MG and IC regions in marmosets.

### Hierarchical structure and two levels of predictive coding of auditory sequences

Studies using the local-global or human language paradigm have provided empirical evidence that temporal processing is hierarchical (Chao et al., 2018; El Karoui et al., 2015; Uhrig et al., 2014), with early sensory regions affected by the recent properties of the input stream and higher-order regions affected by the more complex features during longer time windows within a stimulus context (Bekinschtein et al., 2009; Hasson et al., 2008; Lerner et al., 2011; Wacongne et al., 2011). However, very few studies have examined the neural mechanism of the succession of prediction-error and prediction signals, that is, how low-level or recent events trigger high-level or prolonged signals. Furthermore, most studies using the local-global paradigm test the global violation by combining xx|xY and xY|xx novelties, which, in fact, contain two different types of predictions. In the paradigm in the present study, xx|xY novelty was only involved in the 2^nd^-level signal with the xY sequence as the internal sequence template, and the xY|xx novelty was involved in both 1^st^- and 2^nd^-level signals (the 1^st^-level novelty triggers the 2^nd^-level novelty), when the xx sequence was the internal template. As demonstrated in our hypotheses and neural results, the separation of the global novelty in the classic local-global paradigm was essential for understanding the neural mechanism of the depth of predictive auditory processing and the hierarchical propagation of prediction-error and prediction-updating information.

Finally, we adopted the omission sequence as a deviant to discern prediction signals at the two hierarchies. The neural activations induced by our deviant stimuli (e.g., xY|xx and xx|xY) cannot conclusively confirm the existence of prediction signals, because the novelty responses may act as modulatory signals of bottom-up input. However, the omission responses allowed us to study top-down prediction signal decoupled from bottom-up input signals, as without sensory input, the neural response to omission cannot be explained by any modulation of feedforward propagation and should contain specific information of upcoming predictive information (Demarchi et al., 2019). Thus, omission responses are considered as the proactive prediction and the violation of this prediction. Although detecting omission could happen retrospectively by comparing input and prediction template after the input is processed (Bendixen et al., 2009), our data demonstrated that responses to the omission only occurred after unexpected omissions (in contrast to the expected xxxx omission task), more likely reflecting the specific prediction signal rather than simple novelty detection (Bendixen et al., 2009; Chennu et al., 2016; Fiser et al., 2016). In the xx task, a fixed play rate of five identical tone ensures robust temporal expectations of the next tone. In line with previous works, the responses to rare omissions in the xx task may reflect a temporal prediction of the 5^th^ tone and violation of the global sequence, regardless of the particular identity of the omitted tone. Our observation of x_|xx sequence only in temporal and frontal cortices using both fMRI and ECoG, may thus support that response of prediction signal does not require bottom-up thalamocortical drive (Demarchi et al., 2019). In contrast, the responses of omissions could be found in xY task, only if both the timing and identity of the 5^th^ tone could be predicted, because it was different from the previous four tones (Sanmiguel et al., 2013). The early γ-band increases and fMRI signals in auditory cortex, which elicited by comparing x_|xY with x_|xx sequences, may reflect the unfilled specific expectation for identified stimulus because in both conditions temporal information was predicted. Therefore, since bottom-up inputs cannot account for omission responses, the early γ after following omission onset in x_|xY sequence could be the consequence of 1^st^-level top-down prediction. The correlation we observed between the extended γ and long-lasting β signals in both x_|xx and x_|xY sequences implied the interaction between PE2 caused by global violation and 2^nd^- level prediction. The frontal prediction signal in x_|xY sequence for the update of both PE1 and PE2 on the following sequence and the temporal prediction signal for the update of the PE1, consistently supported the hierarchical top-down predictions. Therefore, in conclusion, our neural results from fMRI and ECoG recordings in marmosets may reveal a complete account of a hierarchical predictive coding framework of auditory sequences, including the prediction-error and prediction-updating information at two hierarchies.

## Materials and Methods

### fMRI

Three adult common marmosets (*Callithrix jacchus*; two males, 350–420 g, 48–60 months) were used in the fMRI experiments. Awake marmoset fMRI training was adopted from Silva et al (Silva et al., 2011). Each marmoset was habituated to a simulated scanning environment over ∼2 months. During training, the marmosets lay in a prone position with their heads immobilized by custom-made helmets. During routine fMRI scanning, the marmosets passively listened to auditory stimuli in a fully awake state without any active task. The auditory stimuli were delivered through a pair of MR-compatible in-ear headphones (Model S14, Sensimetrics) at an average intensity of 75 dB. The protocol of the fMRI study was approved by the Ethical Committee of the Institute of Neuroscience, Chinese Academy of Sciences (no. ION-20180522).

fMRI was performed with a 9.4 T Bruker BioSpec MRI scanner (Bruker, Billerica, MA), with an 8-channel phased array coil for marmoset brain imaging. The BOLD fMRI data were collected using a gradient-echo echo-planer imaging sequence with the sparse acquisition scheme so that a repetition time (TR) consisted of both acquisition time (TA, 1.5s) and silence duration (2s). The auditory stimuli were only played in the silence period to avoid any auditory interference. The acquisition parameters for fMRI data were as follows: TR, 3.5 s; echo time, 15 ms; TA, 1.5 s, field of view, 40×35 mm^2^; matrix, 80×70; voxel size, 0.5×0.5×1 mm^3^; number of slices, 33. A total of 116 volumes were acquired during each run.

### ECoG implantation and recording

Two adult common marmosets (*Callithrix jacchus*; monkey J, male; monkey N, female; 450– 470 g, 42–90 months) were used in the ECoG study. Before the implantation of the ECoG array, the marmosets were familiarized with the experimental settings. The marmosets were then chronically implanted with whole-cortical 96-channel ECoG arrays in the epidural space. The arrays covered the left hemisphere of each marmoset, including most of the frontal, motor, parietal, temporal, and occipital cortices. The details of the surgical procedures for electrode implantation are provided in a previous study (Komatsu et al., 2019). The coordinates of recording electrodes were identified on the basis of the combination of pre-acquired MR images and postoperative computer tomography images.

ECoG signals were acquired using a Grapevine NIP system (Ripple Neuro, Salt Lake City, UT) at a sampling rate of 1 kHz. During the ECoG recordings, the marmosets sat in a sphinx position in an electrically shielded and sound-attenuated chamber with their head fixed. The auditory stimuli were delivered bilaterally by two audio speakers (Fostex, Japan) at a distance of ∼20 cm from the head at an average intensity of 70 dB. All procedures of the ECoG study were conducted in accordance with a protocol approved by the RIKEN Ethical Committee [no. W2020-2-008(2)].

### Auditory paradigm

A classical local-global paradigm was used in the fMRI and ECoG experiments (Fig. 1B). For fMRI, each trial was a sequence of five tones. Each tone lasted 50 ms with an interval of 100 ms. Thus, the total duration of the sequence was 650 ms and the interval of sequence onsets was 3.5 s. At the local level (millisecond timescale), a sequence of five identical tones comprised the local standard (e.g., xxxxx, referred to as “xx”). When the 5^th^ tone was replaced by a distinct tone, the sequence was identified as a local deviant (e.g., xxxxY, referred to as “xY”). In the case of an omission, the sequence consisted of only four identical tones (5^th^ tone omitted; e.g., xxxx). The frequency of the tone x or Y was either 800 or 6,000 Hz. At the global level (second timescale), a series of identical sequences played repeatedly were identified as global standards (e.g., xY). A distinct sequence presented rarely and randomly was identified as the global deviant (e.g., xx).

A given fMRI run started with a resting period of 14 s, following a habitation phase during which the selected global standard sequence was repeatedly presented 20 times to establish a global regularity. In the testing period, three blocks of 25 trials were presented sequentially, each followed by a 14 s rest. The 25 trials included 20 frequent global standards (80%) and five rare global deviants (20%). The global deviants were followed by more than one global standard. Each run lasted 6 min 46 s. In the xx task, the testing blocks were either a mixture of 80% xx trials (xx|xx) and 20% xY trials (xY|xx), or a mixture of 80% xx trials and 20% omission trials (x_|xx). In the xY task, the testing blocks were either a mixture of 80% xY trials (xY|xY) and 20% xx trials (xx|xY), or a mixture of 80% xY trials and 20% omission trials (x_|xY). Each regular local-global session consisted of 3–4 runs of the xx task and 3–4 runs of the xY task, depending on the marmosets’ performance on that day. In the omission local-global experiments, each session similarly consisted of 3–4 runs of the xx task and 3–4 runs of the xY task. The order of the tasks was random, and the frequencies selected for x and Y were balanced.

The duration of each ECoG trial was 850 ms, with each tone lasting 50 ms with an interval of 150 ms. The interval between sequence onsets was 3 s. The frequency used for tone x or Y was either 707 or 4,000 Hz. A separate run consisted of habituation and testing blocks with an omission sequence (xxxx) only. Each session contained 1–2 runs of the xx task with xY|xx or with x_|xx as the deviant, as well as 1–2 runs of the xY task with xx|xY or x_|xY as the deviant.

### fMRI data analysis

The fMRI data were analyzed with Statistical Parametric Mapping (SPM12, http://www.fil.ion.ucl.ac.uk/spm). Time series were slice-time corrected, and the acquired volumes were realigned to the first volume in the series. Data in which the head motion exceeded 0.5 mm or 0.5° were excluded. The realigned images were registered to the NIH marmoset anatomical template (Liu et al., 2018) and then spatially smoothed with a 0.6 mm full-width at half maximum Gaussian kernel.

For first-level analyses, a general linear model was established using experimental conditions (habituation, xx|xx, xY|xx, xY|xY, and xx|xY for regular local-global analysis or habituation, xx|xx, x_|xx, xY|xY, and x_|xY for omission local-global analysis), together with head motion parameters as regressors. At the single-session level, the statistical maps (threshold at *p* < 0.01, uncorrected) generated by all auditory stimuli from a testing period without bilateral activation of auditory cortex were rejected. Therefore, the useful data for regular local-global stimuli comprised 55 sessions (21, 18, and 16 sessions for each of the three marmosets). The useful data for omission local-global stimuli were 48 sessions (18, 11, and 19 sessions for each of the three marmosets).

To determine the pathway of propagation of novelty signals, a group analysis or within-subject analysis of variance (one-way ANOVA) was used for regular and omission local-global experiments, respectively. The single-session contrast images corresponding to each trial type derived from the 1^st^-level analyses were introduced to the 2^nd^-level ANOVA design.

To detect the auditory pathway of marmosets, activation in response to all sound stimuli (except during habituation trials) relative to that at rest was assessed using a threshold of *p* < 0.001 (family-wise error rate corrected) with a cluster size of >50. To determine the regions of 1^st^-level novelty response, the contrast was determined by combining xY|xx – xx|xx and xY|xY – xx|xY using a threshold of *p* < 0.05 (FDR-corrected) and a cluster size of >5. The contrast of 2^nd^-level novelty response was determined using xx|xY – xY|xY (excluding xY|xx – xx|xx). The threshold used was *p* < 0.05 (FDR-corrected) with a cluster size of >10. For both 1^st^- and 2^nd^-level novelty effects in the xx task, the contrast was set as xY|xx – xx|xx (excluding xx|xY – xY|xY) with a threshold of *p* < 0.01 (FDR-corrected) and a cluster size of >10. To investigate the omission effect, the comparisons were x_|xx versus xx|xx using a threshold of *p* < 0.001 (uncorrected) and a cluster size of >10, x_|xY versus xY|xY, and x_|xY versus x_|xx using a threshold of *p* < 0.05 (FDR-corrected) and a cluster size of >10. The significance of x_|xx versus xx|xx was too small to be corrected.

### ECoG data analysis

ECoG data were processed using Fieldtrip toolbox (Oostenveld et al., 2011). Trials were extracted from −1 to 2 s from the onset of the first tone. Each trial was resampled at 500 Hz and bandpass filtered using a bandpass filter order 5 with a window from 1 to 240 Hz. The electrodes with high-frequency noise over 60% of the trials were removed. For all remaining electrodes, trials with abnormal spectra and slow-wave activity were manually rejected (ft_rejectvisual.m). After artifact rejection, a run with fewer than five deviant trials was excluded. The useful data were notch filtered to remove the 50 Hz line noise, re-referenced using a common average reference montage, detrended to remove linear trends, and demeaned to apply baseline correction. The numbers of runs adopted were 23 (monkey J) and 20 (monkey N) for the xx task with xY|xx as the deviant, and 20 (J) and 18 (N) each for the xx task with omissions, xY task with xx|xY, and xY task with omissions.

Time-frequency analysis was conducted using “mtmconvol” in Fieldtrip toolbox (ft_freqanalysis.m). Multitaper time-frequency transformation was based on multiplication in the frequency between 10 and 140 Hz (1 Hz step). The time from 300 ms before the 1^st^ tone to 800 ms after the 5^th^ tone (0.02 s step) was analyzed. Four time windows per frequency were analyzed, and the amount of spectral smoothing through multitapering was 0.5× the frequency. TFR was calculated by comparing deviant and standard trial types with baseline correction. To assess significant differences between TFRs, Monte-Carlo estimates of the significance probabilities were determined from the permutation distribution. Then, a nonparametric cluster-based test was applied for the correction of multiple comparisons over frequency and time (ft_freqstatistics.m). The permutation was performed 1,000 times by shuffling the trial labels “deviant” and “standard.” In each permutation, an independent-sample *t* test was conducted at each frequency and time point. The cluster-level statistics were computed by adopting the sum of the *t* values within each cluster and taking the maximum. The cluster-corrected threshold for significance was set at *p* < 0.05.

To estimate the propagative function among different levels of prediction-error and prediction signals, the contributions of PE1, PE2, and prediction components were measured for each trial. The contribution of each component was achieved from parallel factor analysis using N-way toolbox (http://www.models.life.ku.dk/nwaytoolbox). According to the statistical results of each TFR comparison, the frequency and time domains of different components in each trial type were determined by the significant activity. The trial-by-trial TFR computed from the specific frequency and time domains was projected onto the spatio- spectro-temporal pattern. Then, the values for how much the PE1, PE2 and prediction components contributed to a given trial were obtained. To investigate the relationship among PE1, PE2 and prediction activities during deviant stimuli, the correlation between the contribution of one component derived from the corresponding significant electrodes and the contribution of another within the same type of trial was assessed. For example, the contribution of PE1 component was generated from the electrodes with significant response to 1^st^-level novelty. To observe the role of updating the prediction signal, the correlation between the contribution of the prediction component derived from deviant trials was further evaluated along with the contribution of the PE1 or PE2 component derived from post- deviant trials (which was the standard trial directly after a deviant trial). For the prediction component derived from the electrodes located at frontal cortex that with corresponding significant β-band activity, we calculated the correlation between PE1/PE2 component generated from all electrodes with corresponding significant γ-band activity and the prediction. However, for the prediction component elicited from the significant electrodes located at auditory cortex, we evaluated the correlation between PE1/PE2 component generated from auditory electrodes with corresponding significant γ-band activity and the prediction. If the prediction information was updated, the contributions of the prediction component to deviant trials would significantly correlate to the contributions of the prediction-error component to post-deviant trials. The correlation coefficient from a Pearson correlation with a *p*-value of <0.05 was considered significant.

## Acknowledgements

We thank Yuri Shinomoto and Dr. Takaaki Kaneko for animal care and awake ECoG recordings. We also thank 9.4 T high field small animal MRI platform and Primate physiology research platform from Institute of Neuroscience for data acquisition of awake fMRI.

## Funding

This work was supported by the Strategic Priority Research Program (XDB32070200 and XDB32030100 to L.W., XDBS01030100 to Z.L.), the Pioneer Hundreds of Talents Program from the Chinese Academy of Sciences (to L.W. and Z.L.), the Shanghai Municipal Science and Technology Major Project (2018SHZDZX05 to L.W. and 2018SHZDZX05 to Z.L.), the Instrument Functional Developing Project from the Chinese Academy of Sciences (to L.W.), the National Natural Science Foundation of China (81801354 to Z.L. and 31900797 to Y.J.), the Youth Innovation Promotion Association Chinese Academy of Sciences (to Y.J.), the Brain/MINDS from the Japan Agency for Medical Research and Development (JP20dm0207069 to M.K.) and JSPS KAKENHI (JP19H04993 to M.K.).

## Competing interests

The authors declare no competing financial interests.

---

**Supplementary File 1:** fMRI activation maps of three individual monkeys and ECoG responses of monkey N.

**Figure S1:**
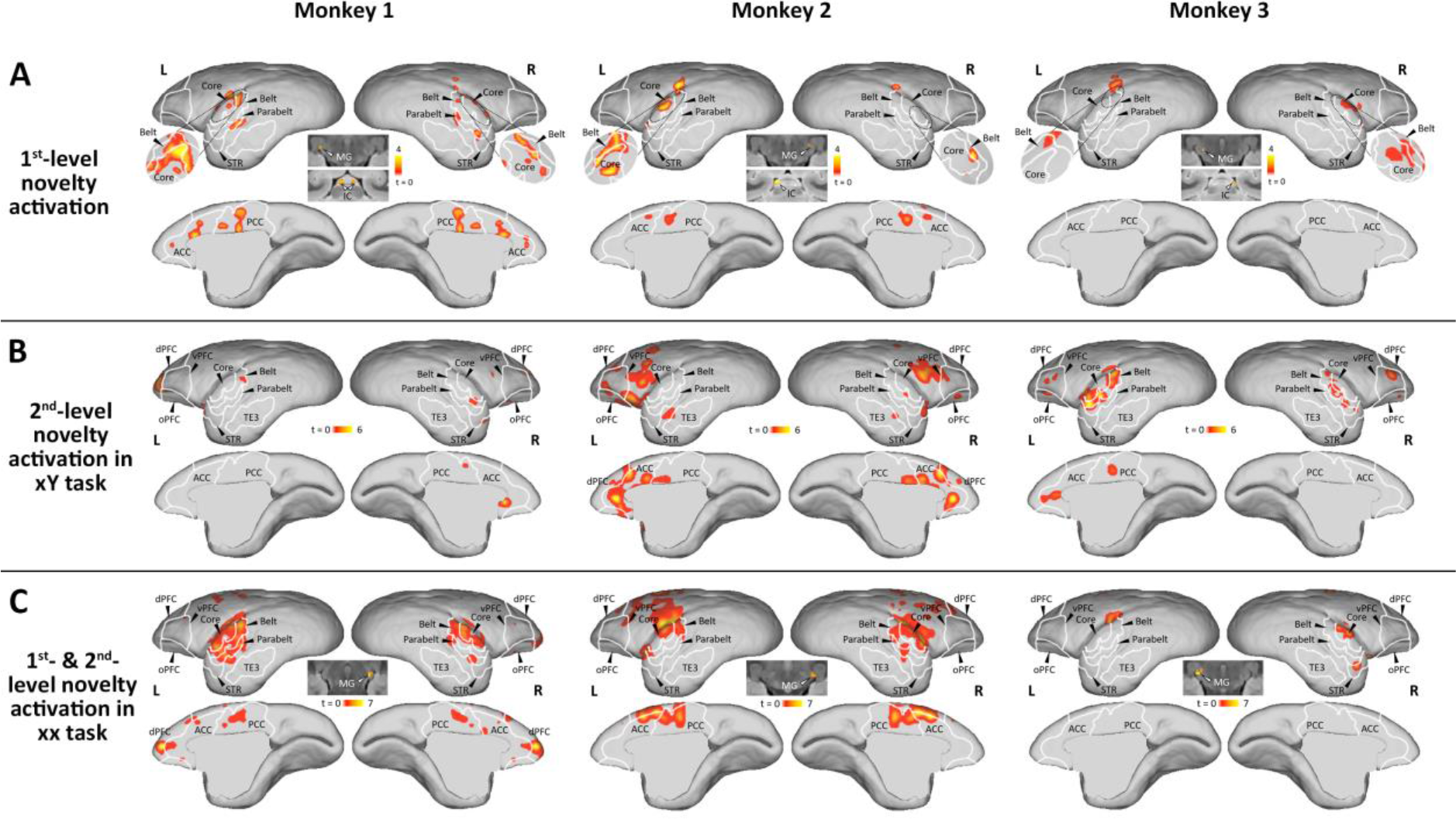
The whole brain fMRI activations to different level novelties in three individual monkeys during regular local-global experiment. **(A)** fMRI activated areas for 1^st^-level novelties (*p* < 0.005, uncorrected, cluster size > 10). **(B)** fMRI activated areas for 2^nd^-level novelties in xY task (*p* < 0.001, uncorrected, cluster size > 10). **(C)** fMRI activated areas for both 1^st^- and 2^nd^-level novelties in xx task (*p* < 0.001, uncorrected, cluster size > 10). The same format is used as in Fig. 2A and 3A.

**Figure S2:**
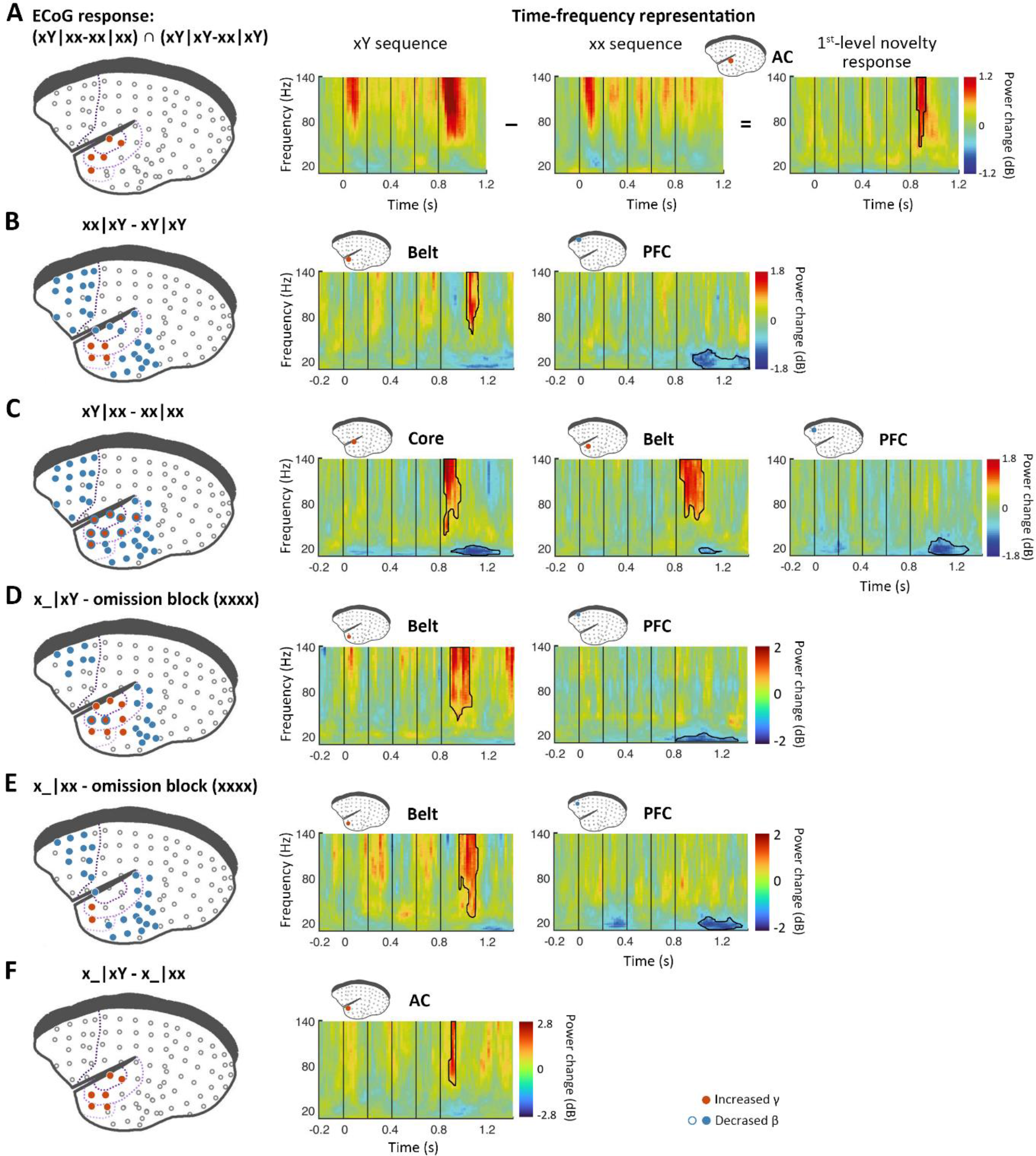
ECoG responses to different types of novelties (in monkey N). **(A)** *Left*, localization of ECoG electrodes with significant responses to 1^st^-level novelties (*p* < 0.05, cluster-corrected). *Right*, Time-frequency representation of representative electrodes. **(B-F)** Significant ECoG responses to 2^nd^-level novelties (B), successive 1^st^- and 2^nd^-level novelties (C), omission of the 5^th^ tone of local deviants (D), omission of the 5^th^ tone of local standards (E), difference between local deviant and local standard omissions (F) (*p* < 0.05, cluster-corrected), with the same format as for panel A.

**Figure S3:**
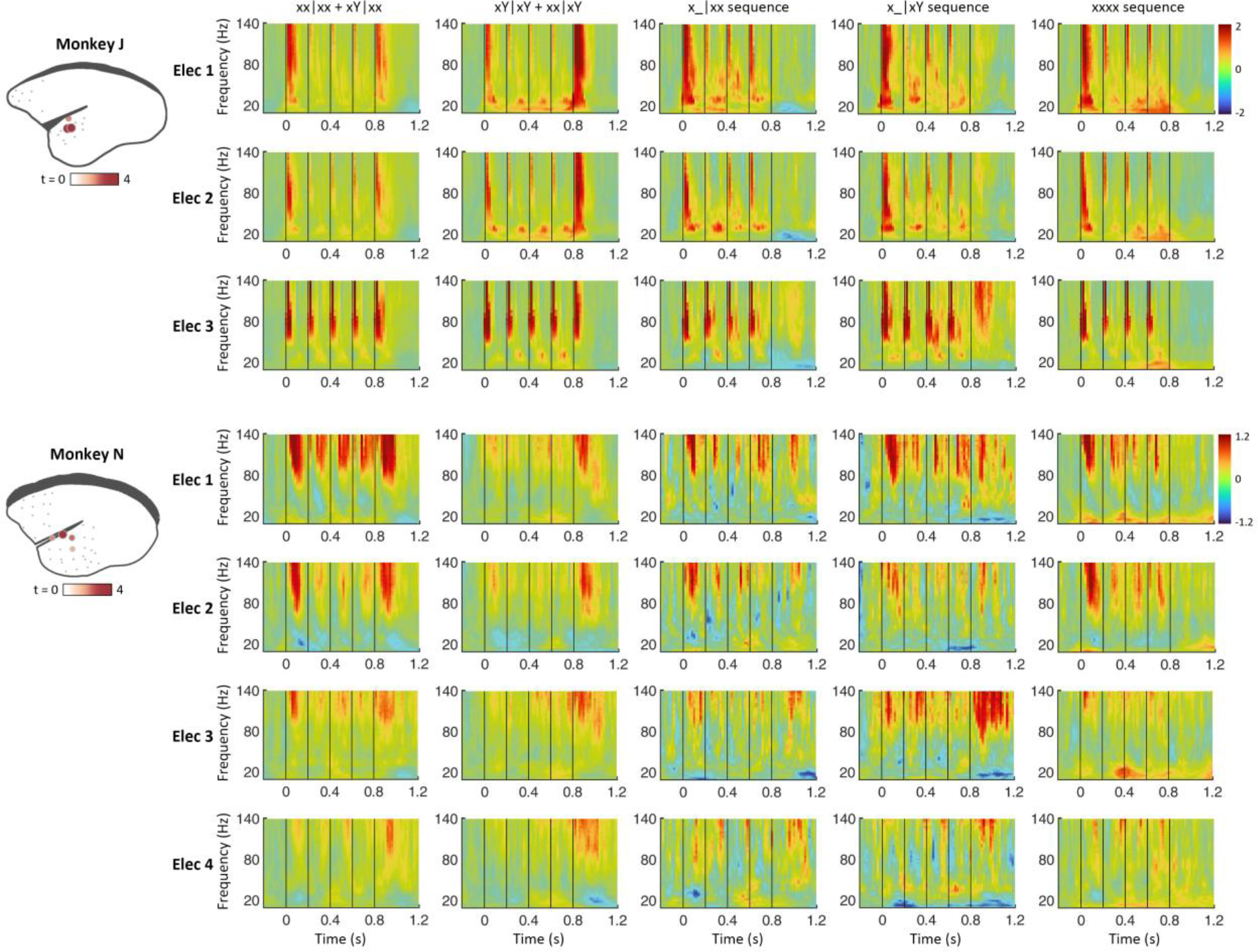
Responses to different sequence types from electrodes with significant auditory response. *Left*: the electrodes showing significant response to auditory stimuli, when compared all auditory sequence with baseline, regardless of the sequence type. The color and size of each dot indicate the level of activation. *Right*: illustration of different type time-frequency representations (TFRs) of the electrodes with significant auditory response. The TFRs are the averaged activation in all sequences in the test period in xx task (xx|xx and xY|xx, column 1), all sequences in the test period in xY task (xY|xY and xx|xY column 2), all x_|xx sequences (column 3), all x_|xY sequences (column 4) and all sequences in omission block (column 5). Electrode (Elec) number indicates the strength order of activation.

**Figure S4:**
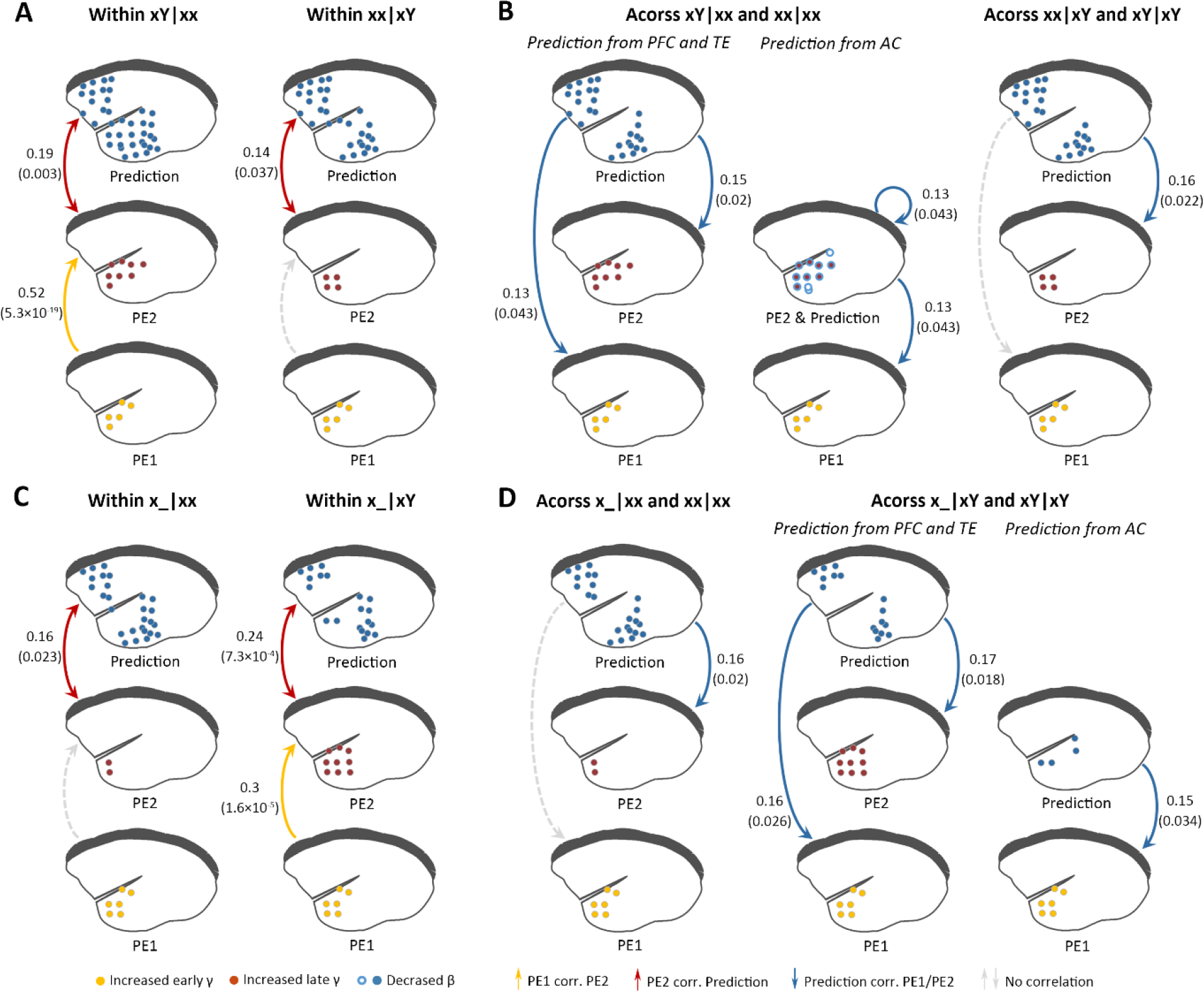
Assessment of propagative roles among different levels of prediction error and prediction signals (Monkey N). **(A)** The functional correlation within xY|xx (left) and within xx|xY (right) trials. **(B)** The functional correlation across xY|xx and subsequent xx|xx (left) trials, and across xx|xY and subsequent xY|xY (right) trials. **(C)** The functional correlation within x_|xx (left) and within x_|xY (right) trials. **(D)** The functional correlation across x_|xx and subsequent xx|xx (left) trials, and across x_|xY and subsequent xY|xY (right) trials. (A-D) with the same format as Fig. 3C and D, Fig. 5C and D.

**Figure S5:**
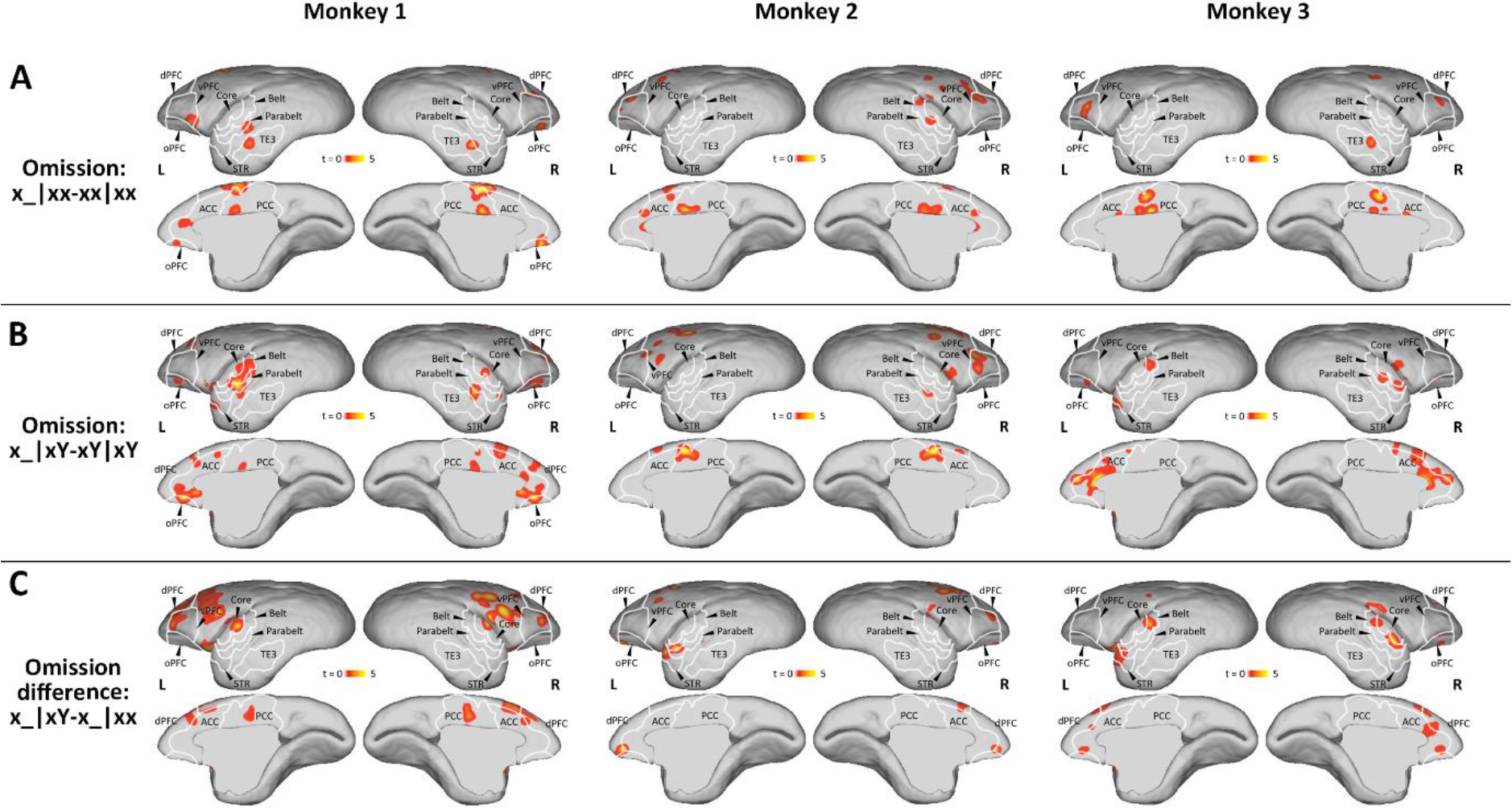
The whole brain fMRI activations to omission of the 5^th^ tone in three individual monkeys. **(A)** fMRI activated areas for omission of the 5^th^ tone of local standards (*p* < 0.005, uncorrected, cluster size > 10). **(B)** fMRI activated areas for omission of the 5^th^ tone of local deviants (*p* < 0.005, uncorrected, cluster size > 10). **(C)** fMRI activated areas for difference between local deviant and local standard omissions (*p* < 0.005, uncorrected, cluster size > 10). The same format is used as in Fig. 4A and 5A.

**Supplementary File 2:** Details of fMRI results for brain regions of intrest.

**Table S1:**
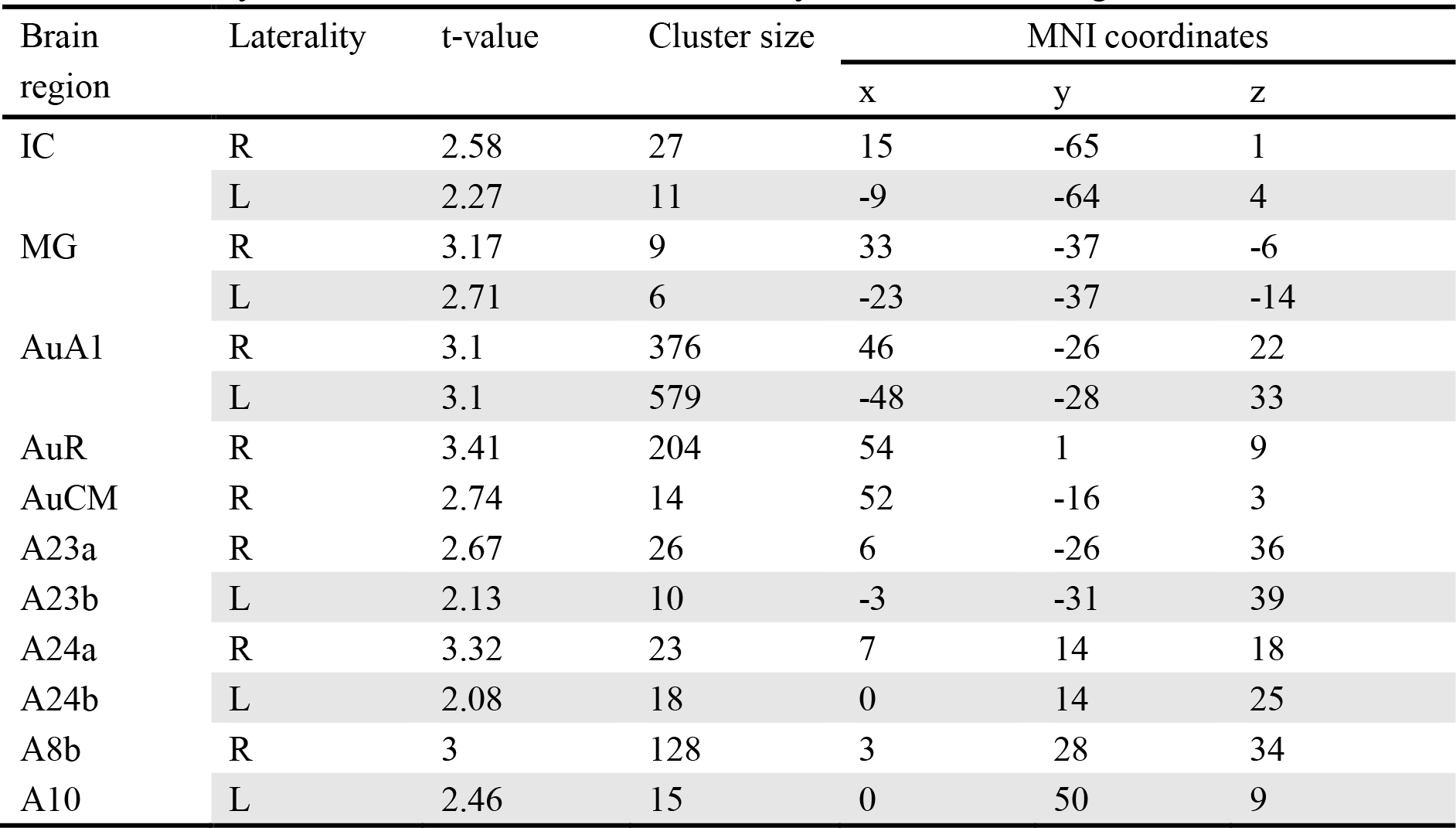
The peak fMRI activations of brain regions for 1^st^-level novelty. The image results were zoomed into human brain size for visualization and using MNI coordinates. IC, inferior colliculus; MG, medial geniculate nucleus; AuA1, primary auditory area; AuR, rostral auditory area; AuCM, caudomedial auditory area. L, left; R, right.

| Brain region | Laterality | t-value | Cluster size | MNI coordinates |  |  |
| --- | --- | --- | --- | --- | --- | --- |
|  |  |  |  | x | y | z |
| IC | R | 2.58 | 27 | 15 | -65 | 1 |
|  | L | 2.27 | 11 | -9 | -64 | 4 |
| MG | R | 3.17 | 9 | 33 | -37 | -6 |
|  | L | 2.71 | 6 | -23 | -37 | -14 |
| AuA1 | R | 3.1 | 376 | 46 | -26 | 22 |
|  | L | 3.1 | 579 | -48 | -28 | 33 |
| AuR | R | 3.41 | 204 | 54 | 1 | 9 |
| AuCM | R | 2.74 | 14 | 52 | -16 | 3 |
| A23a | R | 2.67 | 26 | 6 | -26 | 36 |
| A23b | L | 2.13 | 10 | -3 | -31 | 39 |
| A24a | R | 3.32 | 23 | 7 | 14 | 18 |
| A24b | L | 2.08 | 18 | 0 | 14 | 25 |
| A8b | R | 3 | 128 | 3 | 28 | 34 |
| A10 | L | 2.46 | 15 | 0 | 50 | 9 |

**Table S2:** The peak fMRI activations of brain regions for 2^nd^-level novelty by comparing xx|xY with xY|xY. The image results were zoomed into human brain size for visualization and using MNI coordinates. AuAL, anterolateral auditory area; STR, superior temporal rostral area.

| Brain region | Laterality | t-value | Cluster size | MNI coordinates |  |  |
| --- | --- | --- | --- | --- | --- | --- |
|  |  |  |  | x | y | z |
| AuAL | R | 3.58 | 28 | 61 | -8 | -6 |
|  | L | 4.19 | 78 | -60 | -2 | -11 |
| AuCM | R | 5.72 | 97 | 45 | -13 | 10 |
|  | L | 4.6 | 22 | -54 | -31 | 10 |
| STR | R | 4.72 | 120 | 54 | 7 | -20 |
| A32 | L | 4.5 | 265 | -8 | 34 | 9 |
| A23a | R | 4.45 | 32 | 10 | -5 | 22 |
|  | L | 4.73 | 123 | -8 | -20 | 28 |
| A3b | R | 5.65 | 271 | 21 | -22 | 39 |
| A4ab | L | 4.5 | 46 | -30 | -5 | 36 |
| A8aV | R | 5.29 | 162 | 19 | 41 | 16 |
|  | L | 4.82 | 142 | -27 | 25 | 16 |
| A13 | L | 4.91 | 237 | -11 | 26 | -6 |

**Table S3:** The peak fMRI activations of brain regions for both 1^st^- and 2^nd^-level novelty by comparing xY|xx with xx|xx. The image results were zoomed into human brain size for visualization and using MNI coordinates. AuRT, rostrotemporal auditory area; AuRTM, rostrotemporal medial auditory area; AuRPB, rostral parabelt auditory area; PE, parietal area.

| Brain region | Laterality | t-value | Cluster size | MNI coordinates |  |  |
| --- | --- | --- | --- | --- | --- | --- |
|  |  |  |  | x | y | z |
| MG | R | 5.12 | 44 | 25 | -40 | -11 |
|  | L | 5.64 | 80 | -29 | -38 | -6 |
| AuA1 | L | 6.37 | 1597 | -63 | -14 | 15 |
| AuRT | R | 6.99 | 1258 | 57 | -2 | 0 |
|  | L | 4.3 | 25 | -63 | 4 | -3 |
| AuRTM | L | 5.33 | 118 | -54 | -2 | -8 |
| AuRPB | R | 4.14 | 24 | 60 | -7 | -18 |
| A31 | L | 4.82 | 88 | -8 | -31 | 51 |
| A23b | R | 4 | 16 | 6 | -23 | 37 |
| PE | L | 5.17 | 580 | -17 | -23 | 43 |
| A3a | L | 4.4 | 96 | -36 | -5 | 39 |
| A8b | R | 3.77 | 26 | 9 | 26 | 37 |
| A10 | L | 3.81 | 34 | -12 | 53 | 13 |

**Table S4:** The peak fMRI activations of brain regions for omission of the 5^th^ tone in xx task by comparing x_|xx with xx|xx. The image results were zoomed into human brain size for visualization and using MNI coordinates. TE3, inferior temporal cortex.

| Brain region | Laterality | t-value | Cluster size | MNI coordinates |  |  |
| --- | --- | --- | --- | --- | --- | --- |
|  |  |  |  | x | y | z |
| TE3 | R | 4.09 | 30 | 63 | -28 | -24 |
|  | L | 3.88 | 23 | -65 | -28 | -24 |
| A8aV | R | 3.99 | 46 | 36 | 35 | 21 |
| A47L | L | 3.83 | 13 | -45 | 28 | 1 |
| A23a | R | 3.77 | 17 | 3 | -10 | 27 |
|  | L | 5.12 | 155 | -11 | -16 | 25 |
| A23b | R | 4.23 | 45 | 9 | -20 | 39 |
|  | L | 4.98 | 92 | -5 | -19 | 45 |
| A30 | R | 3.9 | 69 | 9 | -26 | 22 |

**Table S5:** The peak fMRI activations of brain regions for omission of the 5^th^ tone in xY task by comparing x_|xY with xY|xY. The image results were zoomed into human brain size for visualization and using MNI coordinates. AuML, middle lateral auditory area.

| Brain region | Laterality | t-value | Cluster size | MNI coordinates |  |  |
| --- | --- | --- | --- | --- | --- | --- |
|  |  |  |  | x | y | z |
| AuML | R | 4.06 | 19 | 64 | -17 | 0 |
|  | L | 4.84 | 41 | -60 | -26 | 10 |
| A13M | R | 5.26 | 68 | 16 | 28 | -2 |
|  | L | 4.9 | 62 | -23 | 25 | -5 |
| A47M | R | 4.61 | 43 | 30 | 29 | 2 |
| A24d | R | 4.88 | 71 | 15 | -2 | 36 |
| A32 | L | 4.15 | 30 | -11 | 37 | 9 |

**Table S6:**
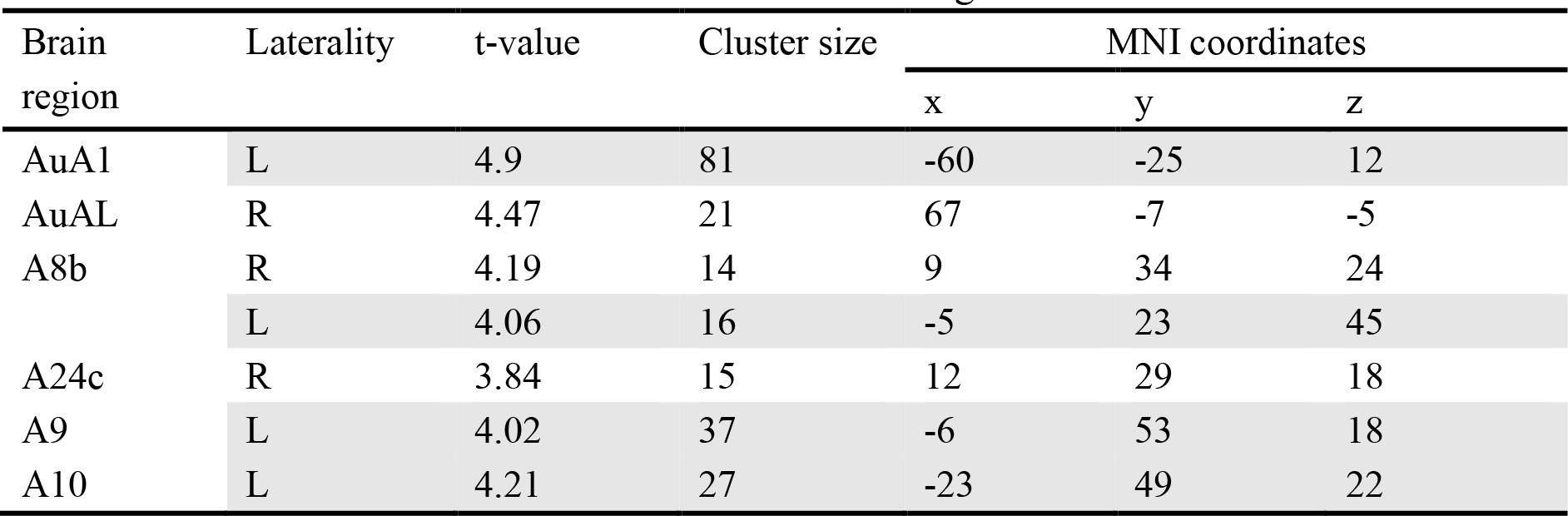
The peak fMRI activations of brain regions for difference between local deviant and local standard omissions by comparing x_|xY with x_|xx. The image results were zoomed into human brain size for visualization and using MNI coordinates.

| Brain region | Laterality | t-value | Cluster size | MNI coordinates |  |  |
| --- | --- | --- | --- | --- | --- | --- |
|  |  |  |  | x | y | z |
| AuA1 | L | 4.9 | 81 | -60 | -25 | 12 |
| AuAL | R | 4.47 | 21 | 67 | -7 | -5 |
| A8b | R | 4.19 | 14 | 9 | 34 | 24 |
|  | L | 4.06 | 16 | -5 | 23 | 45 |
| A24c | R | 3.84 | 15 | 12 | 29 | 18 |
| A9 | L | 4.02 | 37 | -6 | 53 | 18 |
| A10 | L | 4.21 | 27 | -23 | 49 | 22 |

## Notes

### Competing Interest Statement

The authors have declared no competing interest.

